# A Bayesian Generative Model of Vestibular Afferent Neuron Spiking

**DOI:** 10.1101/2020.02.03.933150

**Authors:** Michael Paulin, Kiri Pullar, Larry Hoffman

## Abstract

Using an information criterion to evaluate models fitted to spike train data from chinchilla semicircular canal afferent neurons, we found that the superficially complex functional organization of the canal nerve branch can be accurately quantified in an elegant mathematical model with only three free parameters. Spontaneous spike trains are samples from stationary renewal processes whose interval distributions are Exwald distributions, convolutions of Inverse Gaussian and Exponential distributions. We show that a neuronal membrane compartment is a natural computer for calculating parameter likelihoods given samples from a point process with such a distribution, which may facilitate fast, accurate, efficient Bayesian neural computation for estimating the kinematic state of the head. The model suggests that Bayesian neural computation is an aspect of a more general principle that has driven the evolution of nervous system design, the energy efficiency of biological information processing.

**Significance Statement:** Nervous systems ought to have evolved to be Bayesian, because Bayesian inference allows statistically optimal evidence-based decisions and actions. A variety of circumstantial evidence suggests that animal nervous systems are indeed capable of Bayesian inference, but it is unclear how they could do this. We have identified a simple, accurate generative model of vestibular semicircular canal afferent neuron spike trains. If the brain is a Bayesian observer and a Bayes-optimal decision maker, then the initial stage of processing vestibular information must be to compute the posterior density of head kinematic state given sense data of this form. The model suggests how neurons could do this. Head kinematic state estimation given point-process inertial data is a well-defined dynamical inference problem whose solution formed a foundation for vertebrate brain evolution. The new model provides a foundation for developing realistic, testable spiking neuron models of dynamical state estimation in the vestibulo-cerebellum, and other parts of the Bayesian brain.

## Introduction

The vestibular organs enable agile movement and perceptual acuity by providing the brain with sense data for spatial orientation and postural stability. Among the five sensory epithelia within the mammalian vestibular labyrinth, three semi-circular canal cristae each detect head rotation around a single axis. Dedicated branches of the vestibular nerve transmit Information from each semicircular canal to the brain (Goldberg et al., 2012). Early recordings indicated that the population firing rate within each nerve branch encodes the rate at which the head is turning around the canal axis (Lowenstein & Sand, 1940), but single-unit recordings later revealed systematic, correlated statistical and dynamical heterogeneity within each population (Goldberg & Fernandez, 1971b). The pattern of vestibular afferent neuron behaviour is similar in all vertebrates and has been described using a variety of mathematical models (Paulin & Hoffman, 2019), but remains unexplained. Why is low-dimensional sensory information about head rotation around a single axis distributed across such a large number of channels in parallel? Why are the spike trains so noisy? Why are the statistical and dynamical properties of these neurons so diverse, and why are they correlated? At first sight information transmission in the vestibular nerve seemed simple: Firing rate encodes stimulus strength. But it turns out to be much more complicated than that. Why?

We hypothesized that these questions can be answered by modelling the activity of vestibular sensory afferent neurons as observations for a Bayesian observer, whose goal is to infer what in the world is causing the observations. In this paper we explain how we identified a Bayesian generative model of spontaneous firing in vestibular semi-circular canal afferent neurons, and how it may provide a foundation for modelling neural mechanisms of perception as Bayesian inference.

A Bayesian observer represents relevant states of the environment and themselves using a probability distribution, called the Bayesian posterior distribution. They apply Bayes rule to infer the posterior, the conditional probability distribution of states given what they observe (Gelman et al., 2013; Jaynes & Bretthorst, 2003; Kruschke, 2015). Bayesian inference allows statistically optimal evidence-based decisions and actions (Berger, 1985). This has led to speculation that our nervous systems ought to have evolved to be Bayesian, with selective fitness as an optimization criterion (Doya, 2007; Knill & Pouget, 2004; Knill & Richards, 1996; Kording, 2007; Kording & Wolpert, 2006; Levy, 2006; O’Reilly, Jbabdi, & Behrens, 2012; Ramirez & Marshall, 2017; Yuille & Kersten, 2006). The behaviour of humans and other animals is consistent with this “Bayesian brain” hypothesis (Ostwald et al., 2012; Valone, 2006). However, because Bayesian inference is conditional not only on observations but also on a model of how observations depend on states, and optimality criteria can be arbitrary, it is possible to reverse-engineer a Bayesian explanation for any observed behaviour (Bowers & Davis, 2012; Jones & Love, 2011). Thus realistically modelling neural computation for Bayesian inference, and testing the Bayesian brain hypothesis, requires neurobiological model systems whose performance can be quantified independently and for which observer models can be determined empirically. We suggest that the vestibular system, including the vestibulo-cerebellum, which has long been proposed as a locus of Bayesian neural computation for dynamical estimation of head kinematic state variables (Borah, Young, & Curry, 1988; de Xivry, Coppe, Blohm, & Lefevre, 2013; MacNeilage, Ganesan, & Angelaki, 2008; Paulin, 1989, 1993, 2005; Paulin & Hoffman, 2011; Selva & Oman, 2012; Young, 2011), is suitable for this purpose.

Except in some classical special cases, dynamical Bayesian inference requires a generative model, a model capable of generating simulated observations with the same statistical distribution as the data. Given such a model, sequential random sampling methods can be used to infer the Bayesian posterior density of the model parameters from data (Doucet, De Freitas, & Gordon, 2001). Mathematical parameters of a realistic generative model will map onto to kinematic state variables of the head, the physical parameters of vestibular afferent neuron spike trains. Thus a first step towards a realistic model of Bayesian neural computation for optimal dynamical head kinematic state estimation in the vestibulo-cerebellum is to identify a generative model of vestibular sensory afferent neuron spike trains.

## Methods

All procedures involving animals were approved by the UCLA Chancellor’s Animal Research Committee, and conformed to guidelines mandated in the *NIH Guide for the Care and Use of Laboratory Animals* (National Institutes of Health Publication, revised 2011), and the *Guidelines for the Use of Animals in Neuroscience Research* (Society for Neuroscience).

### Animal preparation

Adult male chinchillas (n=27; body mass 450 – 650 grams) were used in these experiments. They were first anesthetized with isoflurane, after which an intravenous cannula was secured within a jugular vein through which maintenance doses of sodium pentobarbital (0.05cc, 50 mg/cc) were administered. A tracheotomy was performed into which a catheter delivering 100% O_2_ was loosely placed. Heart and respiratory rates, as well as O_2_ saturation levels, were monitored throughout the surgical preparation and recording session. Core body temperature was maintained between 38° - 38.5°C with a custom servo-controlled heater and rectal thermocouple probe. Animals remained physiologically stable throughout the long electrophysiologic recording sessions, which at times lasted longer than 12 hours.

Upon achieving a surgical plane of anesthesia animals were fit into a custom head holder fixed to a turntable. Surgical procedures were similar to those utilized in previous investigations of vestibular afferent electrophysiology (Baird, Desmadryl, Fernandez, & Goldberg, 1988). The right middle ear was exposed by removing the bony cap of the tympanic bulla. The bony ampullae of the superior and horizontal semicircular canals were identified, which provided landmarks to the internal vestibular meatus channelling the superior vestibular nerve between the labyrinth and brainstem. The superior vestibular nerve was exposed at this site, approximately 1 – 2 mm from the landmark ampullae, using fine diamond dental drill bits. Final exposure of the nerve was achieved by gently teasing the epineurium from the nerve with electrolytically sharpened pins.

### Single afferent electrophysiology

Spontaneous discharge epochs from 330 semicircular afferents within the superior vestibular nerve were recorded with high-impedance microelectrodes (40 – 60MΩ) driven by a piezoelectric microdrive. Spontaneous discharge was detected as the electrode approached an afferent, and generally improved with subtle adjustments in electrode position achieved by small manipulations of the microdrive (e.g. small forward and reverse displacements, in addition to gentle tapping of the drive). Upon achieving stable recording, manual turntable displacements were used to identify the epithelium from which the afferent projected. Afferents innervating the horizontal and superior cristae increased their discharge to rotations resulting in utriculofugal and utriculopetal endolymph flow, respectively, and would decrease in discharge in response to turntable rotations in the opposite direction. Afferents projecting to the utricle were generally unresponsive to rotations, or increased their discharge during application of rotations in both directions (centripetal displacements of the otolithic membrane concomitant with rotation in either direction). These afferents were excluded from the present dataset.

### Spiketrain analysis and model fitting

#### Data acquisition, Summary Statistics and Exploratory Analysis

Single-unit spike times were acquired in 20-second records with 300μs resolution, and imported into MATLAB as arrays of interspike interval (ISI) lengths in seconds. Plots of spike time data and ISIs were visually inspected to identify trends, discontinuities and outliers indicating possible miss-triggering during data acquisition. We tested for serial correlation in interval length using a Wald-Wolfowitz runs test (MATLAB function *runstest*). Records with detectable artefacts or non-stationarity were removed, leaving 306 (of the initial 330) selected records for further analysis and modelling.

Mean 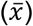, standard deviation (*s*), coefficient of variation 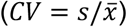 and Pearson’s moment of skewness (*γ* = *E*[(*x* − *µ*)^3^]/*σ*^3^) were computed for the intervals in each spiketrain, using MATLAB functions *mean*, *std* and *skewness*. Standard deviations of interval length for the most regular units in our sample are comparable to the resolution of spike time data acquisition (300us). Because of this, estimates of CV and skewness for very regular units may be less reliable than estimates for irregular units. CV is a scale-invariant measure of variability. It is near zero for highly regular spike trains, near 1 for completely random or Poisson-like activity, and becomes larger than 1 for clumped or bursting activity. By convention, neurons whose CV falls in the lowest 1/3 of a sample of vestibular afferents are deemed “regular”, neurons whose CV falls in the largest 1/3 are deemed “irregular”, while neurons with intermediate CV are deemed “intermediate” (Goldberg & Fernandez, 1971b).

#### Candidate Models

The selected records are observations from a stationary renewal process (no correlation or trends in interval length over time), which can be modelled as sequences of samples from a fixed-parameter probability distribution of interval lengths. This is a complete model because the event times themselves, up to an arbitrary start time, can be recovered from the sequence of intervals between them. Since interval lengths must be positive and can have arbitrary length, candidate models must be probability density functions *f*(*t*; *α*) defined on *t* > 0 with parameters *α*.

Previous studies have shown a consistent pattern of ISI distributions in vestibular afferent spike trains. ISI distributions of the most regular afferents have narrow distributions which are nearly symmetrical and approximately Gaussian, with standard deviations much smaller than mean interval length (*σ ≪ µ*). A Gaussian with *σ* ≪ *µ* > 0 has essentially no probability mass below zero and can be treated as a density on *t* > 0. ISI distributions of more irregular neurons tend to be more right-skewed with larger CVs, while interval distributions of the most irregular neurons resemble exponential distributions, with standard deviation similar to mean interval length (CV=1). Differences between the most regular and the most irregular neurons are so great that it has often been suggested that there are distinct populations within the nerve, but there is a continuum of behaviour between these extremes (Paulin & Hoffman, 2019). Suitable candidate models therefore are positive-valued, continuously-parameterized probability densities whose shape transforms continuously between limiting cases resembling Gaussian and Exponential distributions.

Our candidate models fall into three groups. The first group (1.1-1.5 below) were all initially derived as models of simple physical processes that are at least somewhat analogous to the canonical “noisy integrate-and-fire” model of a stochastic neuron (ref), and have all been applied previously to model spiking statistics of neurons, including vestibular semicircular canal afferent neurons (refs). This group contains the Weibull, Log-normal, Erlang (Integer Gamma), Birnbaum-Saunders (cumulative damage) and Inverse Gaussian or Wald distributions. They are available in the MATLAB Statistics Toolbox.

##### 1.1 Weibull

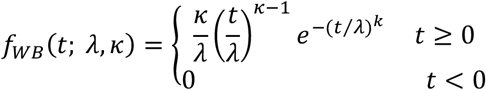

is the distribution of intervals between events when event rate is proportional to a power of the waiting time since the last event. This is a birth-death model with “aging”. When *κ* = 1 (constant event rate) the Weibull reduces to an Exponential distribution.

##### 1.2 Log-normal

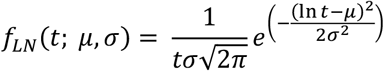

is the distribution of outcomes of a growth process involving multiplicative interactions among many small random effects. Multiplicative interactions are additive on a log scale, so the log of the outcome has a Gaussian or normal distribution because of the Central Limit Theorem.

##### 1.3 Erlang

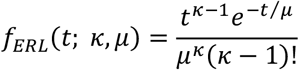

where the *shape parameter*, *κ*, is a positive integer and the *scale parameter*, *µ*, is a positive real number, is the distribution of waiting times for *κ* events in a Poisson process when the average waiting time is *µ* (such that the average waiting time in the underlying Poisson process is *µ*/*κ*). When *κ*=1 the Erlang reduces to an Exponential distribution, the waiting time distribution for events in a Poisson process. This has been a popular model of neuronal firing variability, including for vestibular afferent neurons, because of its flexible shape which resembles empirical interval distributions, and because it has a simple mechanistic interpretation as the waiting time for the accumulation of quantal events occurring at random times to reach a threshold (Lansky, Sacerdote, & Zucca, 2016; Shimokawa, Koyama, & Shinomoto, 2010).

##### 1.4 Birnbaum-Saunders

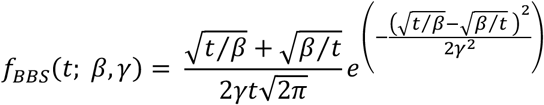

is the distribution of waiting time for the accumulation of events with a Gaussian distribution of amplitudes occurring at random times to reach a threshold. It is also known as the Cumulative Damage distribution because of its application to modelling time-to-failure of a system subjected to impacts with random magnitudes occurring at random times. It is a physically plausible model of time to threshold for a neuron receiving EPSPs with Gaussian amplitudes, which fits spike train data from real neurons and biophysically realistic computational neural models (Leiva et al., 2015).

##### 1.5 *Inverse Gaussian* or Wald

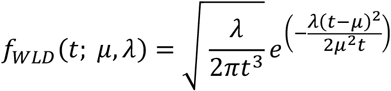

is the distribution of waiting times for Gaussian noise with mean 1/*µ* with and variance 1/*λ* to integrate to a threshold at 1. It models the first passage time (time to hit a barrier) of a drift-diffusion process, i.e. Brownian motion in constant flow (Chhikara & Folks, 1989; Folks & Chhikara, 1978).

As discussed in the Results section, a second group of candidate models was constructed by adding a fixed latency (time offset) parameter to some of the candidates in Group 1. This group contains Erlang, Wald and Birnbaum-Saunders distributions, each with an additional time-shift parameter, *τ*.

##### 2.1 Offset Erlang

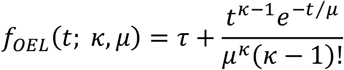

##### 2.2 Offset Wald

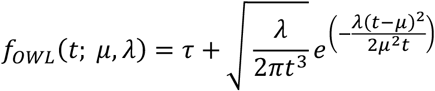

##### 2.3 Offset Birnbaum-Saunders

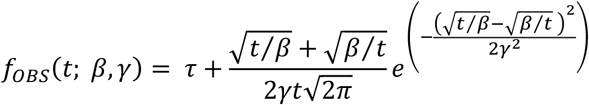

For reasons discussed in Results, a third group of models was constructed by replacing the constant offset parameter *τ* in the Group 2 models with an Exponentially-distributed random time offset having mean *τ*. In each case this creates a new random variable as the sum of two random variables, whose distribution is the convolution of the distributions of the components.

##### 3.1 Exerlang

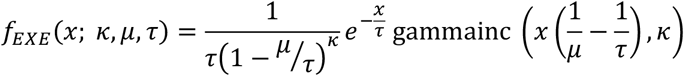

This expression for the convolution of an Exponential distribution and an Erlang distribution was obtained using *Mathematica* (Wolfram Research, Illinois, USA). *gammainc* is the MATLAB incomplete gamma function, a MATLAB built-in special function. The incomplete gamma function is defined slightly differently in MATLAB and *Mathematica*, so the result derived by *Mathematica* requires adjustment to obtain the formula given above.

##### 3.2 Exwald

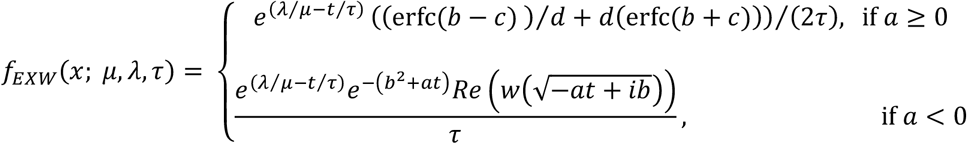

where 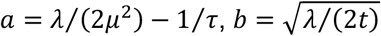 and 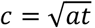. erfc is the complementary error function, *w* is the Fadeeva scaled complex complementary error function (Abramowitz & Stegun, 1964), 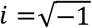 and *Re*(*z*) is the real part of the complex number *z*. This expression was modified from formulae given by Schwarz (ref), by setting the barrier distance/threshold level parameter in the Wald component of Schwarz’s derivation to 1 and scaling the other parameters accordingly. We found that this expression can be numerically unstable when *λ ≪ µ* (diffusion negligible compared to drift) or *τ ≪ µ* (Exponential component negligible compared to Wald component). In the former case we reduced the Wald drift-diffusion component to a pure drift, approximating the Exwald using an Exponential distribution with fixed time offset, *µ*. In the latter case we removed the Exponential component, approximating the Exwald using only the Wald component. None of our data were fitted by models with parameters in regions of parameter space where these approximations were applied, but it was necessary to include these approximations to prevent numerical instability when the fitting algorithm explores the parameter space before converging.

##### 3.3 Exgaussian

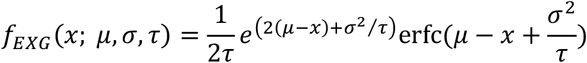

This expression for the convolution of a Gaussian distribution with mean *µ* and variance *σ*^2^ and an Exponential distribution with mean interval parameter *τ* was derived analytically using *Mathematica* (Wolfram Research, Illinois, USA). In this expression, 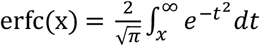, is the complementary error function, a MATLAB built-in Special Function.

#### Fitting Criterion

Given an observed probability distribution, *p*(*t*), and a model *q*(*t*), the Kullback-Liebler divergence from *q*(*t*) to *p*(*t*), also known as entropy of *p*(*t*) relative to *q*(*t*), is

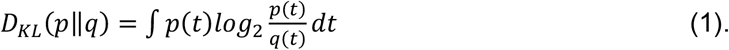

It measures the bits of information lost when *q*(*t*) is used to approximate the empirical distribution, *p*(*t*). Given a set of candidate models, minimum *D*_*KL*_ identifies the candidate that minimizes the expected information in future observations, given what has been observed (Jaynes & Bretthorst, 2003; Kullback & Leibler, 1951; Paulin & Hoffman, 2001).

Given *N* observations *t*_1_, *t*_2_, ⋯, *t*_*N*_, the empirical distribution can be represented as a normalized frequency histogram, with probability *p*_*k*_ = *n*_*k*_/*N* in the *κ* th bin, where *n*_*k*_ is the number of observations in the *k* th bin. Assuming that *q*(*t*) ≈ *q*_*k*_ is constant in the kth bin, the expression for *D*_*KL*_ reduces to a sum,

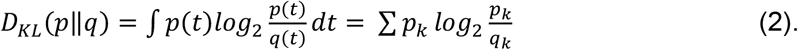

If each bin is very narrow and contains at most one observation then *q*(*t*) = *q*_*k*_ and the normalized histogram reduces to a particle model, with probability *p*(*t*_*k*_) = 1/*N* at the observed points *t*_*k*_ and zero elsewhere. In that case the expression for *D*_*KL*_ reduces to

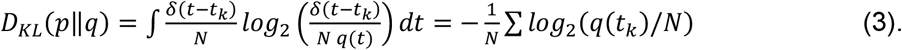

Thus

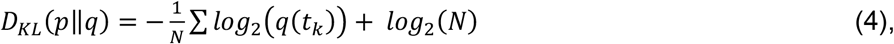

is negative log-likelihood with a logarithmic penalty on sample size.

Since the sample size is fixed in each record, fitting a model by minimum *D*_*KL*_is equivalent to fitting a model by maximum likelihood for any given neuron. However, across neurons KLD scales the log-likelihood by the entropy of the empirical distribution, giving a measure of model performance which is independent of differences in variability of spike time data from different neurons. For example, regular neurons have narrow ISI distributions with high probability densities, and generate more spikes during the 20-second recording period because they fire faster. As a result, the likelihood for any given model is generally larger for more regular neurons, and using maximum likelihood would bias section in favour of candidates that are better at fitting regular neurons. *D*_*KL*_ avoids this problem. Having said that, we found that using maximum likelihood as a model-selection criterion leads to qualitatively similar results as using *D*_*KL*_, and does not change our conclusions.

#### Model Fitting

Models were fitted using the MATLAB function fminseachbnd 1.4.0 (D’Errico, 1965), which implements the Nelder-Mead simplex algorithm (Nelder & Mead, 1965) with constraints. The constraints were applied to prevent the algorithm from stepping outside the region of parameter space in which a model is defined (e.g. negative mean interval length), which would produce meaningless results and/or numerical instability.

#### Analysis of Fitted Models

Candidate models have at most 3 parameters meaning that fitted parameters for each neuron can be visualized as a point in 3D, and parameters fitted to all records form a cloud in 3D space. The cloud of points fitted to our data is roughly ellipsoidal in log-log axes. We computed the major axes of this ellipsoid using the *pca* function in the MATLAB Statistics Toolbox. We computed the convex hull of parameter estimates in 2D projections (the smallest polygon enclosing all points) using the MATLAB built-in function convhull. We used the first principal component axis to generate curves in parameter space showing the predicted value of a parameter given some other parameter. For example, to show how a model parameter *α* relates to the summary statistic 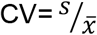, we find parameters on the first principal component axis corresponding to a model with this CV. Simple closed expressions can be found in all cases, i.e. it is not necessary to use numerical optimization/search procedures to compute these curves.

## Results

### Summary Statistics

Figure 1(a) is a scatterplot of conventional ISI summary statistics, mean and coefficient of variation. It shows the heterogeneity of spontaneous discharge characteristics, and the tendency for neurons with shorter mean intervals (higher firing rates) to have more regular firing patterns. The average mean interval is 16.9 ms (± 13.0ms) and the average CV is 0.17 (± 0.22). This plot closely resembles scatterplots of mean ISI vs CV in previous reports of vestibular afferent neuron spiking activity (e.g. Baird et al. (1988) figure 1; Honrubia, Hoffman, Sitko, and Schwartz (1989) figure 6b; Goldberg (2000) figure 3A; Hullar et al. (2005) figure 1). The scatterplot shows the wide variation in mean interval length and CV with no indication of distinct groups within the population.

**Figure 1:**
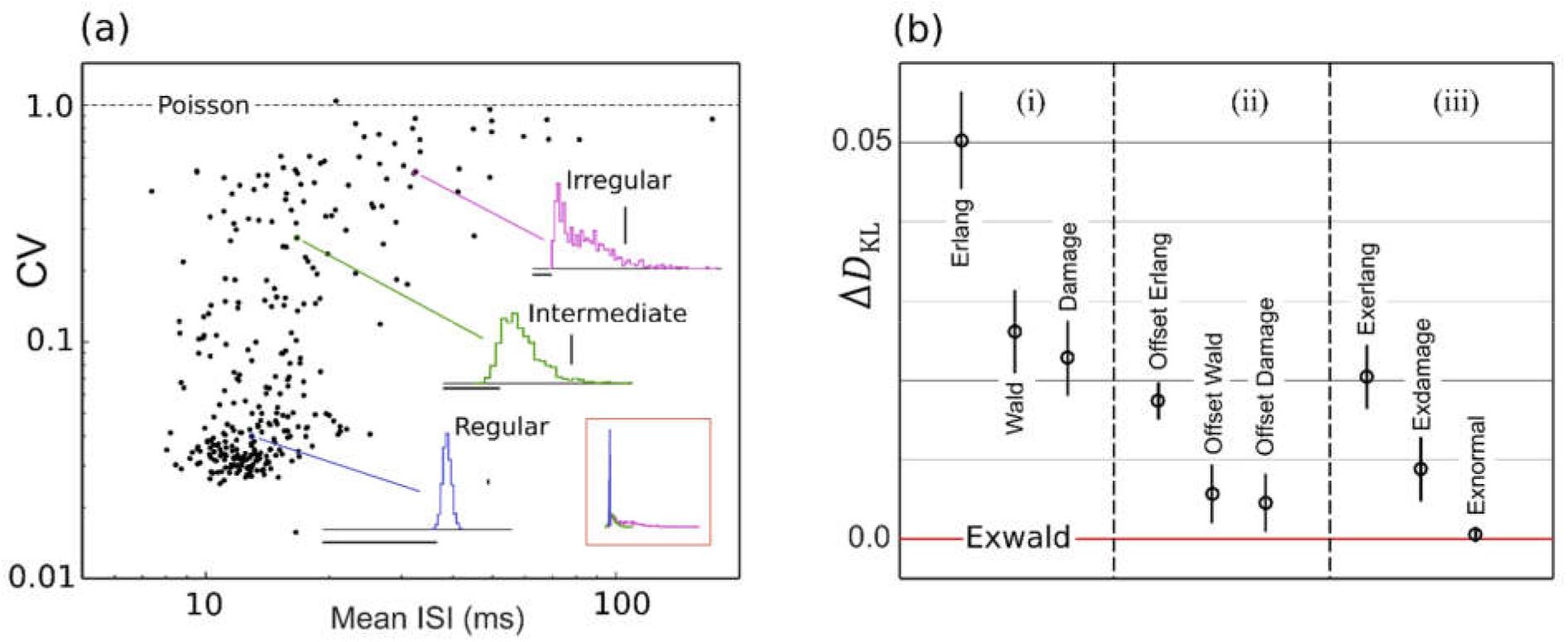
**(a)** Mean inter-spike interval length vs coefficient of variation (CV) of spontaneous activity. Normalized inter-spike interval (ISI) histograms shown for a regular, intermediate and an irregular afferent. Aspect ratios are adjusted to show differences in shape between the distributions. Horizontal scale bars are 12ms. Vertical scale bars represent relative frequency 0.05. Inset (lower right) shows the three histograms overlaid with a common aspect ratio. **(b)** Average Kullback-Liebler divergence (information loss in bits) for candidate models relative to Exwald, the loss-minimizing candidate. Group (i) Random walk models; (ii) Fixed time-offset random walk models; (iii) Exponential random offset random walk models. Vertical bars are standard errors of means.

ISI histograms for three selected afferents are overlaid on the scatterplot. They closely resemble ISI distributions previously reported in vestibular afferents in various species (Paulin & Hoffman, 2019). The inset shows these three distributions plotted on common axes. This illustrates that while mean and CV reveal substantial diversity in spontaneous behaviour of these neurons, these descriptive statistics fail to characterise the shapes of ISI distributions and the large, systematic shape changes across the population. Regular afferents, with faster mean firing rates tend to have narrow, approximately Gaussian ISI distributions, while irregular, slower-firing afferents tend to have positively skewed ISI distributions. The most irregular afferents, with CVs near 1, have ISI distributions that resemble right-shifted or left-censored Exponential distributions. Exponential interval distributions are characteristic of Poisson processes, for which the average time between events is fixed but event times are random (Haight, 1967; Landolt & Correia, 1978). These distributions have the unique property that removing intervals shorter than some specified duration (left-censoring) is equivalent to right-shifting the distribution by that duration.

### Model Fitting

Initial candidate models were continuous probability distributions defined on positive intervals: Weibull, Log-normal, Erlang or Integer Gamma, Inverse Gaussian or Wald, and Birnbaum-Saunders or Cumulative Damage Distribution (See methods). For brevity, we refer to the Birnbaum-Saunders/Cumulative Damage Distribution as the Damage distribution. These candidates were selected because they possess the requisite property of having Gaussian-like shapes in some subregion of their parameter space and Exponential-like shapes in some other subregion. Weibull and Lognormal candidates were quickly eliminated because of large qualitative discrepancies between the shapes of data and model distributions, evident by inspection of plots.

The remaining candidates, Erlang, Wald and Damage distributions, all seem capable of generating the shapes of the empirical interval distributions. In addition, they are all waiting time distributions for random counting or integrating processes to reach a threshold, and can be interpreted in terms of simple models of physical mechanisms that underlie neuronal spiking. All have previously been proposed as models of neuronal spiking variability (See Methods). Each of these distributions has two free parameters.

The relative goodness of fit for these three models is shown in the left column of figure 1(b). The vertical axis in this figure (Δ*D*_*KL*_) is the mean difference between Kullback-Leibler Divergence from model to data for each model, and the Kullback-Leibler Divergence from model to data for the model that was ultimately identified as the best model according to the minimum Kullback-Leibler criterion (See *Methods*). Error bars represent the standard error of mean Δ*D*_*KL*_. According to the minimum Δ*D*_*KL*_ criterion, the Damage distribution is the best of these candidates, followed by the Wald and the Erlang.

Inspection of plots of best-fitting models overlaid on the empirical interval distributions showed that in many cases a fitted model deviated systematically from the data, while manual adjustment of parameters indicated that the model should be capable of fitting the shape of the empirical distribution much more accurately than it did. We hypothesized that this may be because the parameters of these models do not affect shape and location independently. A change in either parameter is generally accompanied by a change in the location (mean) and the shape of the distribution. Because the Kullback-Liebler criterion harshly penalizes models that assign negligible probability to values that are actually observed, minimum Δ*D*_*KL*_ favours spreading probability mass across all observations (i.e. getting the location right) over matching the shape of the empirical distribution, when it is not possible to do both.

We tested this hypothesis by adding a time-offset parameter, allowing each model distribution to shift arbitrarily along the time axis independently of shape changes. The second panel in figure 1(b) shows that this additional offset parameter improves the fit of each model. Visual inspection of plots showed that all three offset models can accurately locate and match the shapes of the empirical distributions. The performance improvement due to the additional free parameter is similar for each model, so that their ranking remains the same. The offset Damage model is the best, followed by the offset Wald and offset Erlang.

Introducing a time offset parameter confirmed that there is (at least) a degree of freedom missing in each of the group 1 statistical models. However, a pure time offset in a model of neuronal spiking is implausible, not simply because it would imply the existence of a biophysical clock mechanism capable of producing precisely-timed intervals of different lengths in different neurons, but because some of the fitted time offset parameters in the group 2 models are negative. This would imply that in some neurons the clock must trigger the counting/integrating process that generates a spike at a precise time before the preceding spike. This would violate causality.

The simplest way to extend the group 1 models in a way that adds a degree of freedom in location is to include a Poisson process in series. A Poisson process has only one parameter, the mean interval length, and has an Exponential distribution of interval lengths (ref). The effect of adding an Exponentially-distributed random delay term to each of the Erlang, Damage and Wald models is shown in the third panel of figure 1(b). This term improves the fit of all three group 1 models. As might be expected, since the time-offset models fit quite precisely and the Poisson series element must introduce a shape change in addition to a time offset, the Poisson element doesn’t improve the fit of the Erlang or Damage models as much as a pure time offset does. Surprisingly, however, it improves the fit of the Wald model by even more than a pure time offset does. Evidently a series Poisson process not only provides an additional degree of freedom allowing the Wald distribution to locate itself over the probability mass of the data, it improves the ability of the Wald distribution to match the shape of the empirical distribution when it gets there.

An Exponential distribution in series with a Wald distribution is called an Exwald distribution (Schwarz, 2001, 2002). Analogously, we refer to the Exponentially-extended Erlang and Damage distributions the Exerlang and Exdamage distributions respectively.

Wald components of fitted Exwald models consistently resemble narrow Gaussians with small positive skewness. Positive skew in an empirical distribution is invariably fitted by increasing the interval parameter of the Poisson component of the model, not by altering the skew of the Wald component. This raises the possibility that positive skew in empirical ISI distributions can be explained entirely by the Poisson component of an Exwald model.

We tested this possibility by adding an Exponential-Gaussian series model to the candidate set. This model is labelled Exnormal in the third panel of figure 1(b). It fits almost as well as the Exwald model on average. The relatively small standard error shows that the Exnormal model fits the data uniformly almost as well as the Exwald model does.

Figure 2 shows Exwald models fitted to ISI histograms for a regular, an intermediate and an irregular unit. These are the same example units shown in figure 1(a). Components of the intermediate model, for which the decomposition is easiest to see, are labelled. All neurons, not just these three examples, have a refractory period of 10-12ms during which the probability of spiking is essentially zero. The refractory period appears to be determined by the Wald component, while the extent of the tail, corresponding to spiking irregularity, appears to be determined by the Exponential or Poisson component. Shape and location parameters of the Wald components are similar for all three neurons, while the interval parameter of the Poisson component is larger for more irregular neurons.

**Figure 2:**
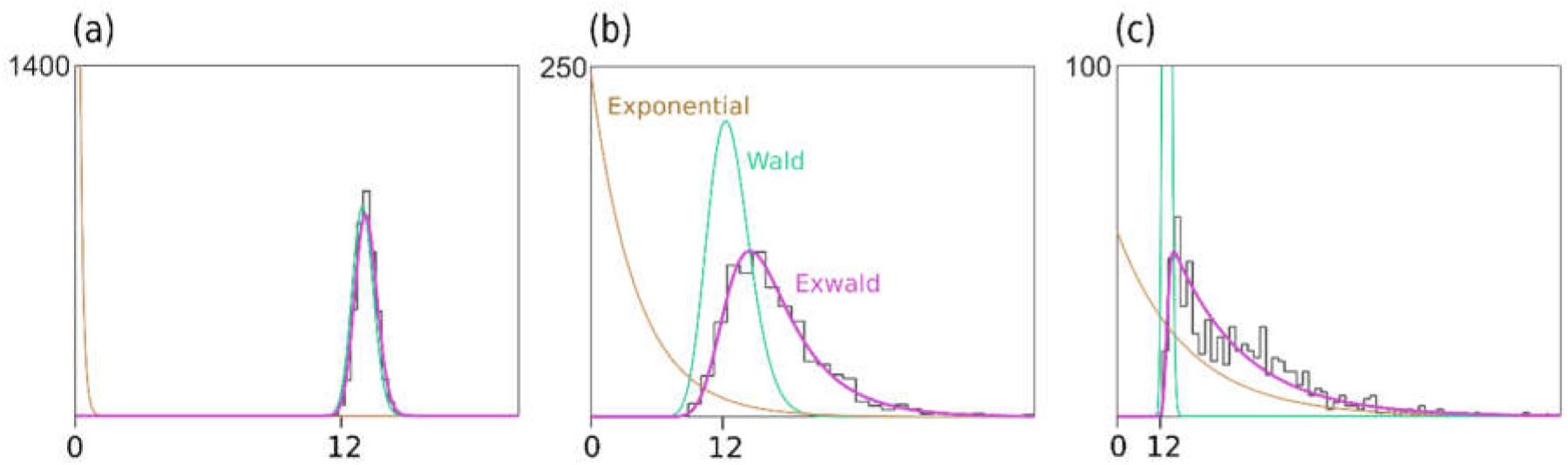
Fitted Exwald models overlaid on empirical ISI histograms for **(a)** a regular **(b)** an intermediate and **(c)** an irregular neuron. The Exwald (pink) decomposes into an Exponential (brown) and a Wald (teal) distribution. Scales are different on each axis because of the large differences in the shapes of the empirical distributions (See inset, figure 1(a)).

### Analysis of the Exwald model

The Exwald is the distribution of intervals generated by an Inverse Gaussian process in series with a Poisson process. Each sample from the Exwald is the sum of a sample from the Wald component and a sample from the Exponential component. It has three parameters: *µ* and *λ*, which are the mean interval and shape parameters of the Wald distribution, and *τ*, which is the parameter of the Exponential interval distribution of the Poisson process. The parameters are all positive quantities with dimensions of time, reported here in milliseconds.

Figure 3 shows the result of principal component analysis (PCA) of Exwald model parameters. The fitted parameters form a flattened, elongated ellipsoidal cloud of points when plotted on log-log axes in 3D. PCA was used to find the major axes of an ellipsoid fitted to this cloud. Panels a-c show the parameter cloud and the principal component axes projected into the three coordinate planes of the parameter space.

**Figure 3:**
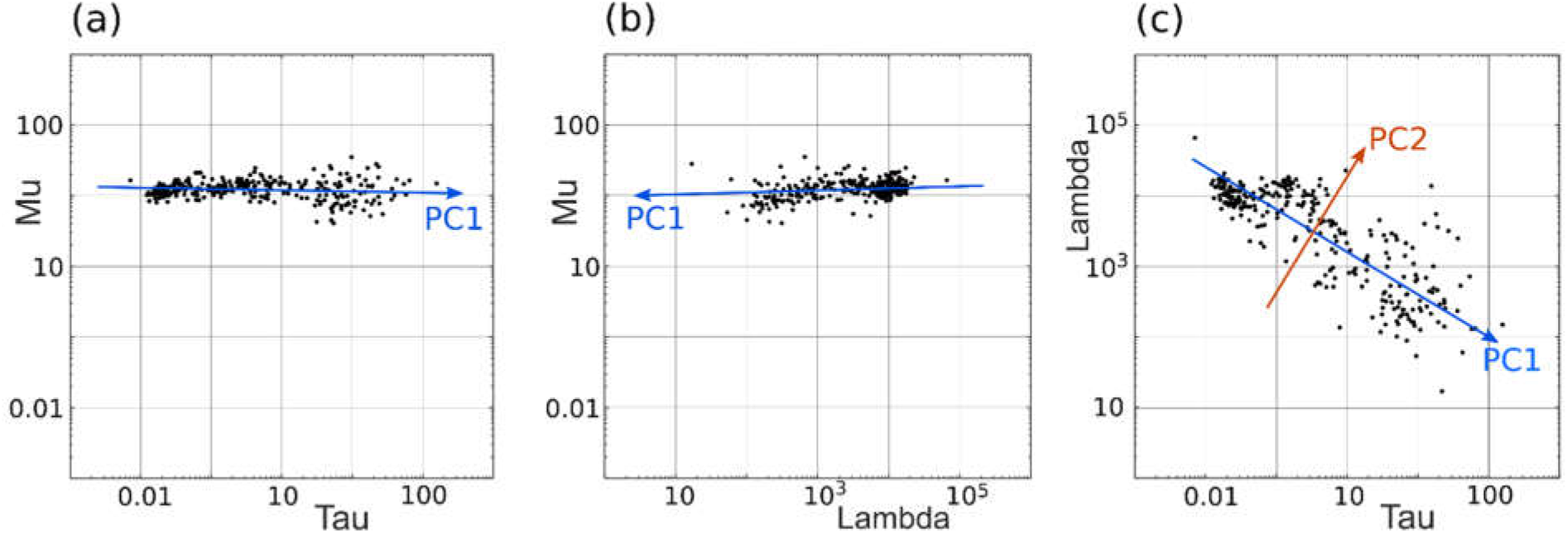
Principal component analysis of Exwald model parameters. **(a-c)** show the 3D cloud of fitted parameters projected into each of the coordinate planes on log-log axes. The aspect ratio is the same on all axes, such that each grid unit represents a tenfold change in magnitude for any of the parameters. Almost all of the variability is in λ and τ, which both vary over several orders of magnitude, while μ is similar, averaging 12.7ms, in all afferents.

Panels (a) and (b) show that the first principal component axis (PC1), the major axis of the parameter distribution, is almost parallel to the *τ* - *λ* plane, with values of *µ* clustered around the mean value of 12.7ms. *τ* varies over roughly 4 orders of magnitude while *λ* varies over roughly 2 orders of magnitude.

Panel (c) shows that most of the variation among parameters, and correspondingly most of the differences between interval distributions, can be explained by only two parameters, *τ* and *λ*. PC1 has a slope near −0.5 in the *τ* - *λ* plane. A slope of −0.5 on log-log axes would indicate an inverse square relationship between these parameters, 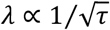.

### Relationship between Exwald model parameters and conventional summary statistics

Figure 4 shows how the parameters of fitted Exwald models are related to the conventional summary statistics that have historically been used to describe the statistical diversity of vestibular afferent firing patterns, mean ISI and CV. The curve in figure 4(a) shows the Exwald model-predicted CV for parameters on the first principal component axis corresponding to a model with the specified mean ISI. It is a projection of PC1 from *log*(*τ*) − *log*(*λ*) parameter space into mean ISI - CV parameter space. It shows that PC1 predicts the nonlinear relationship between mean ISI and CV. Similarly, the curves in figure 4(b) and 4(c) show that *τ* is a good predictor of CV and mean ISI. These plots show that *τ* characterises not only the change in mean and variability of ISI distributions over the population, but also the systematic change in shape of the distributions. For small values of *τ*, 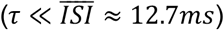, interval length is largely determined by the Wald component, while for large values of *τ*, 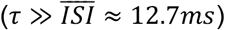, interval length is largely determined by the Poisson component. Thus *τ* by itself characterises the universally observed pattern in which vestibular afferent neurons show a continuous diversity of statistical behaviour from rapidly firing, regular neurons whose interval distributions resemble narrow Gaussians to slowly firing, irregular neurons whose interval distributions resemble right-shifted or (equivalently) left-censored Exponentials.

**Figure 4:**
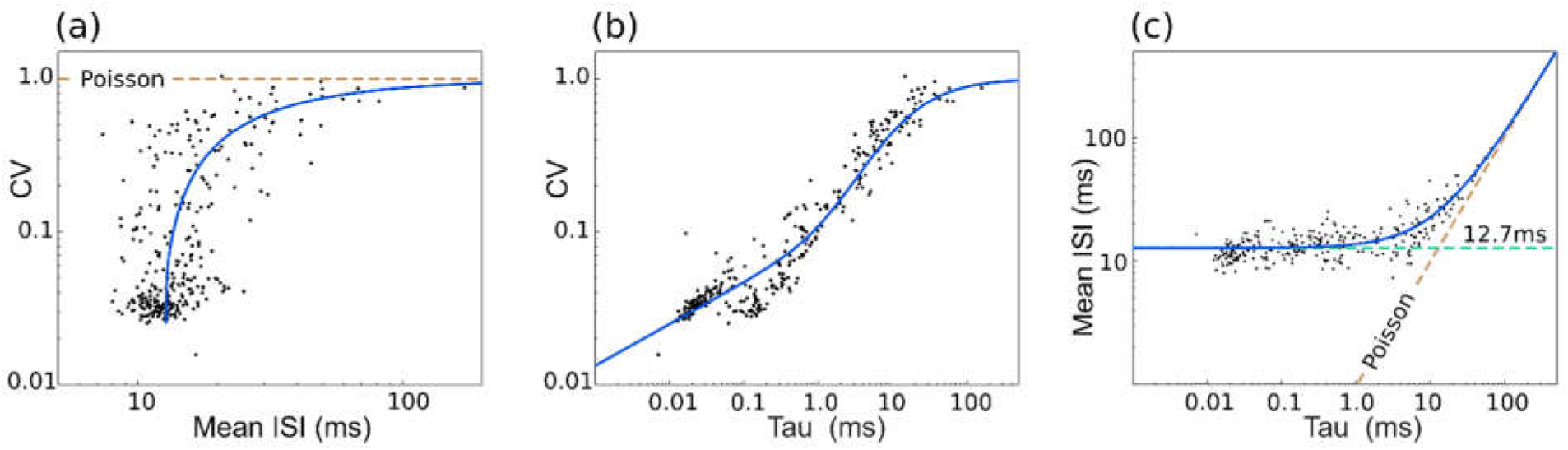
Relationship between parameters λ and τ of the Exwald model and the conventional summary statistics of spontaneous activity, mean inter-spike interval and CV. **(a)** Scatterplot of mean ISI vs CV (c.f. figure 1(a)). Curve shows CV of the Exwald model on PC1 with a given mean ISI. **(b)** Scatterplot of τ vs CV. Curve shows CV of the Exwald model on PC1 for a given τ. **(c)** Scatterplot of τ vs mean ISI. Curve shows the mean ISI of the Exwald model on PC1 for a given τ.

### Distribution of ExWald model shapes in model parameter space

Figure 5 is a map of ISI distributions in *log*(*τ*) − *log*(*λ*) space. Parameter values fitted to data (the same as in figure 3(c)) are plotted here as blue discs. The first two principal component axes are shown. The dashed line is the convex hull, the smallest polygon enclosing all of the fitted parameter points. Shapes of Exwald model interval distributions are drawn on a grid in PC1-PC2 coordinates, aligned with the *log*(*τ*) − *log*(*λ*) axes. For each distribution, t=0 is plotted at the grid point and the time scales are all the same. The vertical (probability density) axes are scaled so that all distributions have the same height on the plot. In reality the distributions for the most regular neurons (upper left of the map) are very much taller than the distributions for the most irregular neurons (lower right). The inset (lower left) shows the true shapes of five distributions spaced along PC1.

**Figure 5:**
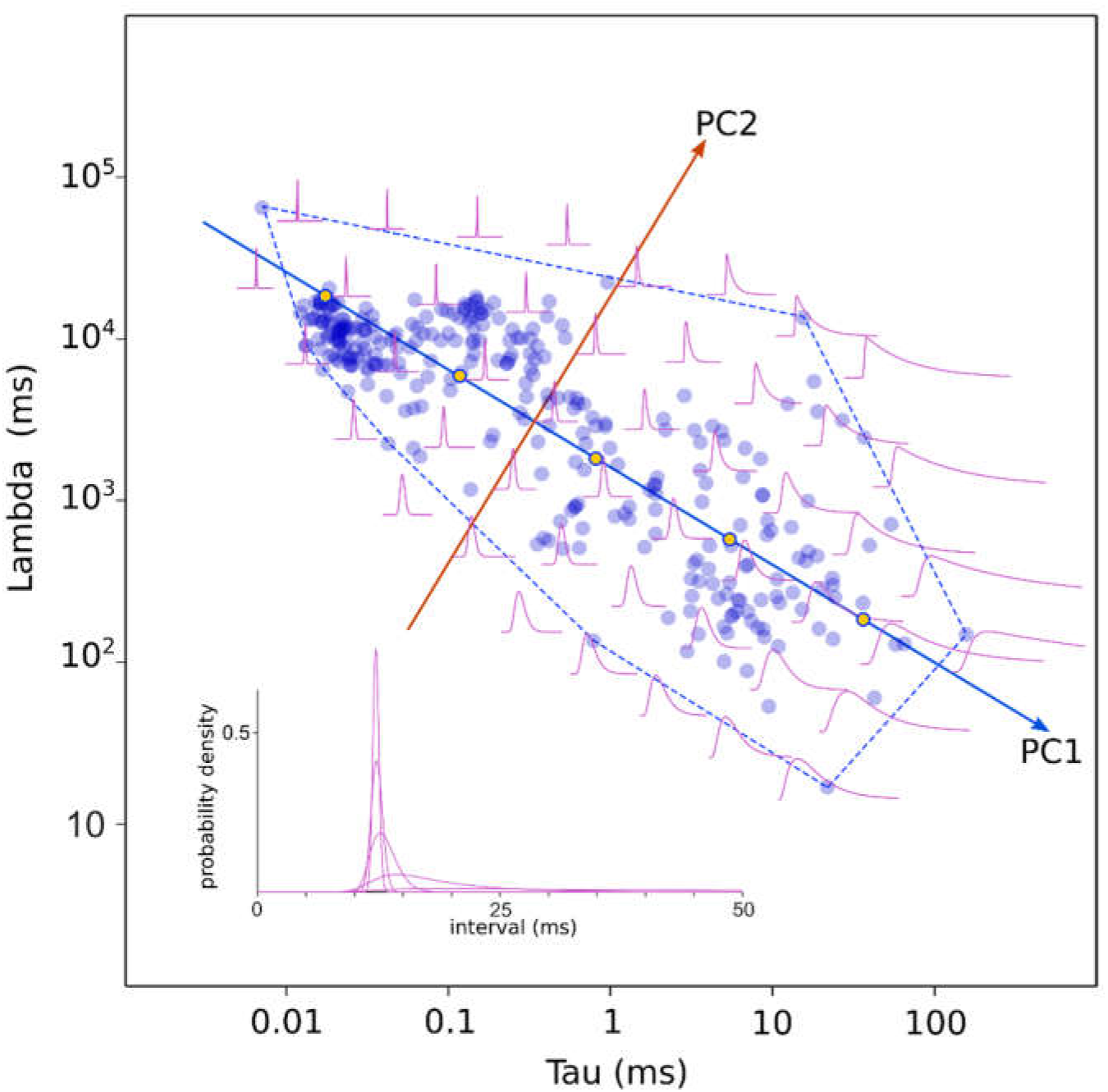
Map of ISI distributions in Exwald parameter space. Scatterplot of fitted parameters in the τ-λ plane (blue discs), with principal component axes projected onto the plane. Dashed blue line is the convex hull of data points. Scaled Exwald models are drawn on a grid. Inset (lower left) shows the true proportions for five models along the first principal component axis. Their parameters are indicated by the five markers along PC1.

This figure indicates that the first principal component measures ISI variability, which is strongly predictable by *τ* (c.f. figure 4(b)), The second principal component measures variability of the refractory period, which is strongly predicted by *λ*, as evidenced by the increasingly steep onset of spiking probability after a refractory period, in the PC2 direction.

According to the principal components analysis, 91.7% of parameter variance is explained by PC1, 7.6% is explained by PC2 and 0.7% by PC3. This suggests that differences in the statistical properties of afferents are mostly controlled by changes in a single degree of freedom in the underlying physical process(es), and are almost entirely, if not entirely, controlled by changes in at most two degrees of freedom.

### Neural Computation for Bayesian Inference Given Exwald Data

The form of the Exwald model suggests an elegant natural mechanism that neurons could employ in inferring the posterior density of stimulus parameters given samples from a point process with Exwald interval statistics. Sequential or dynamical Bayesian inference entails computing the likelihood function for the parameter(s) given the most recent observation, multiplying this by the probability inferred from previous observations (the prior probability) at each parameter value, then (re)normalizing to obtain a function that integrates to 1 over the parameter space (Doucet et al., 2001).

The most recent observation for a stationary renewal process at any time, the latest available information, is the elapsed time since the most recent event. Many models have been proposed to explain multiplicative gain or sensitivity adjustments and normalization of activity levels across neural populations (Bastian, 1986; Beck, Latham, & Pouget, 2011; Capaday, 2002; Carandini & Heeger, 2012; Eliasmith & Martens, 2011; Louie, Khaw, & Glimcher, 2013; Mejias, Payeur, Selin, Maler, & Longtin, 2014; Nelson, 1994; Olsen, Bhandawat, & Wilson, 2010; Silver, 2010), and we will not consider possible mechanisms for these operations in the vestibular system beyond noting that it is widely accepted that neurons are capable of such computations. The key additional computational capability that neurons would require to implement dynamical Bayesian inference in the vestibular system is the ability to compute parameter likelihoods given the elapsed time since the most recent event.

The likelihood function for the parameters *µ*, *λ*, and *τ* of an Exwald process given elapsed time *t* since the most recent event is, by definition, the probability of observing an interval of length *t* if the parameters are *µ*, *λ*, and *τ*. The Exwald distribution is a convolution of an Inverse Gaussian and an Exponential,

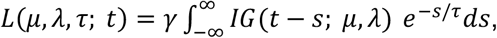

where *γ* is an arbitrary constant (because a scaled likelihood function is a likelihood function). For Bayesian inference given spike train data from a vestibular afferent neuron, (some) central neurons must be able to compute *L*(*µ, λ, τ*; *t*) given *t* > 0.

The electrical response of a neuronal membrane compartment to a transient depolarizing current input with waveform *I*(*t*; *µ, λ*) is the convolution of the exponential impulse response of the membrane compartment with the input waveform,

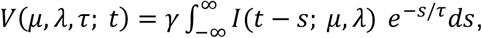

where *V*(*t*) is voltage referenced to resting membrane potential, *τ* is the electrical time constant of the membrane compartment, and *γ* is an arbitrary constant (because current and voltage can be measured in arbitrary units) (Bower & Beeman, 1998; Koch, 1999). Thus a neuronal compartment with membrane time constant *τ*_0_ and a synapse whose EPSP shape is controlled by two parameters, *µ* and *λ*, with values *µ*_0_ and *λ*_0_, is a natural analog computer that instantaneously computes the likelihood for the parameters (*µ, λ, τ*) at the point (*µ*_0_, *λ*_0_, *τ*_0_), at time t after the preceding synaptic input. By postulating that EPSP waveforms *I*(*t*) mimic the shape of Inverse Gaussian distributions *IG*(*t*) - which, on the face of it, they do - neuronal compartments could compute parameter likelihoods from point process data with Exwald interval distributions.

It may be non-trivial to arrange neurons capable of these basic operations (amplification, normalization and evaluating likelihoods) into circuitry capable of inferring the Bayesian posterior density given vestibular afferent neuron spike trains, and will not attempt here to show how it may be done. However, the remarkable isomorphism between equation 1, representing an abstract probability computation which is fundamental for dynamical Bayesian inference, and equation 2, representing a physical model of the electrical behaviour of a neuron, is worth mentioning, because it shows that mechanisms capable of implementing all of the mathematical operations required for dynamical Bayesian inference occur naturally in neurons, and would be available to be co-opted by evolution if there was selection pressure on nervous systems to be Bayesian.

## Discussion

The diversity of vestibular afferent neuron firing behaviour has been characterised in the past using the coefficient of variation (CV) of inter-spike intervals as a signature that predicts other statistical, dynamical and anatomical characteristics of these neurons (Goldberg, 2000; Goldberg et al., 2012). Using an information-theoretic model selection procedure, we found that spontaneous activity patterns of chinchilla semicircular canal afferent neurons can be accurately modelled using a simple, three-parameter model. The first principal component axis of fitted parameters lies almost parallel to the *τ − λ* plane and in log-transformed axes has slope close to -½ in that plane, indicating that *τ* and *λ* are related by a power law that is approximately an inverse square law, 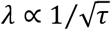. Because there is a unique point on the first principal component axis for any given *τ*, more than 90% of parameter variation among neurons can be explained by *τ* alone. Adding a second parameter, *λ*, accounts for more than 99% of parameter variation. The third parameter, *µ*, contributes less than 1% of parameter variation and is essentially constant for all neurons.

The Exwald parameters, *µ, λ*, and *τ*, can be used to compute the conventional summary statistics of spontaneous activity, mean ISI, CV and skewness, if needed. Because CV is a noisy invertible function of *τ* (figure 4(b)), *τ* by itself should be a good predictor of statistical, dynamical and anatomical properties of vestibular afferent neurons. The Exwald parameters appear to characterise the diversity of spontaneous activity at least as well as the conventional summary statistics do, but in addition they accurately describe the interval distributions themselves.

Because event times can be recovered from the intervals between them, the Exwald model provides a complete stochastic process model of spontaneous activity in these neurons. It can be regarded as a descriptive or phenomenological model whose parameters supersede the conventional summary statistics. Its efficacy as a descriptive model raises the question of whether it is, or may lead to, an explanatory model of vestibular afferent neuron behaviour. Because CV is computable from parameters of an Exwald model and CV is correlated with dynamical response parameters of vestibular afferent neurons, the Exwald model may characterise more than just spontaneous activity patterns of these neurons.

Mechanosensory hair cells and related acousticolateralis receptor cells can transduce signals with power levels smaller than thermal noise power in the transduction mechanisms (Denk & Webb, 1988; Denk, Webb, & Hudspeth, 1986; Markin & Hudspeth, 1995), and such signals are perceptible (Bialek, 1987; Devries, 1948; Torre, Ashmore, Lamb, & Menini, 1995). This implies that stochasticity in spontaneous activity is driven by thermal noise in transduction, synaptic and spike-generating mechanisms. Spontaneous firing is, however, a laboratory artefact, imposed by clamping an animal’s head so that it cannot move. Under natural conditions the head is always moving, and the ecological function of “spontaneous” firing is to provide high acuity sense data for postural stability, compensatory reflexes and acuity of other senses when the animal is not actively moving its head. A completely motionless head is not natural, but it is the limiting case of an ecologically important state. Generalized fluctuation-dissipation theorems then imply that the response to intrinsic thermal noise characterises the system’s dynamical responses to small stimuli (Dinis, Martin, Barral, Prost, & Joanny, 2012; Marconi, Puglisi, Rondoni, & Vulpiani, 2008; Prost, Joanny, & Parrondo, 2009). It follows that an Exwald model fitted to the spontaneous interval distribution of a neuron should be able to predict the neuron’s dynamical responses, at least during small head movements.

When the average firing rate of a vestibular afferent neuron is held at a constant level above its spontaneous rate by applying prolonged unidirectional acceleration, the variability of interval length as measured by CV also increases (Fernandez & Goldberg, 1976; Goldberg, 2000; Goldberg & Fernandez, 1971a). The change in CV as a function of mean interval is approximately a power law 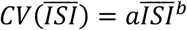 with exponent *b* = 1/2 (Goldberg & Fernandez, 1980; Paulin & Hoffman, 2019). Since CV is the square root of variance divided by the mean, this relationship implies that the variance of interval length scales with the cube of mean interval length, *σ*^2^ ∝ *µ*^3^. This scaling law is a unique signature of an Inverse Gaussian or Wald distribution (Chhikara & Folks, 1989). Thus a simple auxiliary assumption - that vestibular stimulation alters the parameter *µ* - can extend the Exwald model to explain the statistics vestibular afferent neuron responses under constant stimulation.

Instantaneous firing rates of semicircular canal afferents responding to broad-band, naturalistic head motion exhibit a simple, fractional order dynamical relationship to head angular velocity, of the form 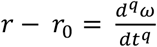, where 0 < *q* < 1 (Paulin & Hoffman, 1999; Paulin & Hoffman, 2019). This suggests that the Exwald model of spontaneous activity may be extended to a stochastic dynamical model by making its parameters depend on head angular velocity in this manner. Further investigation and testing is required in order to determine if and how the Exwald model might be extended to describe the dynamics and statistics of spiking beyond the quasistatic state of small head movements.

The Inverse Gaussian or Wald is the distribution of time taken for Gaussian white noise with mean *r* = *α*/*µ* and power *σ*^2^ = *α*/*λ*^2^ to integrate to a threshold at *α* (ref). We set *α* = 1 without loss of generality (equivalent to choosing units in which *α* = 1). A very simple neural model can explain why the distribution of interval lengths for a stochastic spiking neuron might contain a Wald distribution: If *r* is the mean rate of depolarization, then intervals generated by an integrate-and-fire neuron which resets to zero membrane potential after each spike will have a Wald distribution. An Exwald is the distribution of the sum of samples from a Wald and an Exponential distribution, hinting that vestibular afferent neurons spikes may be generated by a Poisson process in series with a noisy integrate-and-fire process. Poisson distributions occur as limiting cases in many stochastic process models, analogous to the way that Gaussian distributions occur as limiting cases when independent observations are combined (Arratia, Goldstein, & Gordon, 1990; Chen, 1975). Poisson data can be generated simply by threshold triggering in a stationary noise process (Basano & Ottonello, 1975). For example, spontaneous thermal-noise driven opening times of sensory receptor channels are Poisson-distributed, with exponential interval distributions (Sigg, 2014; Smith, 2002).

The existence of very simple mechanisms that can produce events with Inverse Gaussian and Exponential interval distributions then suggests a very simple possible explanation for the superficial statistical complexity of vestibular afferent neuron behaviour. Exwald distributions could have evolved because combining processes with Inverse Gaussian and Exponential interval statistics in series is a simple, feasible way to translate microscopic, low-power stochastic molecular transduction events into high-power electrochemical events carrying the same information but which can be transmitted rapidly over macroscopic distances to the brain (Sterling & Laughlin, 2015). In other words, evolution found a simple way to transmit information from mechanoreceptors to the brain using existing mechanisms, and this happened to produce spike trains with Exwald interval distributions.

However, the molecular mechanisms that mediate signal transmission from transduction in receptor hair cells to spiking in vestibular afferent neurons are prodigiously complex (Glowatzki, Grant, & Fuchs, 2008; Hudspeth, 1983; McPherson, 2018; Vollrath, Kwan, & Corey, 2007). Natural selection appears to have put a great deal of effort into constructing intricate mechanisms that transduce tiny deflections of hair cell cilia into electrical signals, and amplify the transduced signal into spiking events in afferent neurons. The pathway from transduction to spiking is evidently not a simple juxtaposition of two simple molecular mechanisms. On the contrary, it comprises a byzantine conglomeration of structures that collectively behave *as if* this were the case. That the net behaviour of such complex machinery can be accurately modelled in such a mathematically elegant way suggests that the machinery must have been selected to produce this behaviour. That is, there must be some selective advantage in transmitting vestibular information to the brain using spike trains with Exwald-distributed intervals, or some similar distribution.

As discussed in the introduction, there are several ubiquitous characteristics of vestibular afferent neuron behaviour requiring explanation. As noted above, noisiness or stochasticity in afferent spike trains can be explained by specialization to detect and transmit small signals. Spontaneous activity in afferents is driven by thermal noise in molecular mechanisms, which are amplified to produce spike train stochasticity because the peripheral vestibular system has evolved to gather information about signals that are small compared to thermal noise, and amplify them into spike trains (Denk & Webb, 1989, 1992; Denk et al., 1986; Torre et al., 1995; van Netten & Kros, 2000).

The distribution of low-dimensional signals across thousands of afferents can be explained by selection for energy efficiency. The energy cost of spiking neurons, which is dominated by the cost of spiking (Aiello & Bach-y-Rita, 2000; Cohen, 2005; Niven, 2016; Yu & Yu, 2017), is a major constraint on nervous system evolution (Hasenstaub, Otte, Callaway, & Sejnowski, 2010; Laughlin, 2001; Lewis, Gilmour, Moorhead, Perry, & Markham, 2014; Niven & Laughlin, 2008; Sterling & Laughlin, 2015). Because spiking neurons are so energetically expensive, there is strong selection pressure for neurons to maximize channel capacity per unit energy cost. Sterling and Laughlin (2015) suggest that the performance of neural communication and computation should be measured in bits per second per Watt, or bits per Joule, rather than channel capacity, or bits per second, which has been the conventional measure of performance in communication and information processing systems. Using bits per Joule as a proxy for the evolutionary fitness of nervous systems can explain many features of nervous system structure and function (Sterling & Laughlin, 2015).

Spike trains become prohibitively expensive at high average firing rates because when an action potential occurs within a few milliseconds of another, overlapping sodium and potassium ion fluxes consume energy without changing the membrane potential. Efficient neural computation and communication requires average firing rates below about 100s^−1^ (Goldberg, Sripati, & Andreou, 2003; Hasenstaub et al., 2010; Levy & Baxter, 1996). As illustrated in figure 2, semicircular canal afferents have refractory periods in the order of 10ms, and mean interval duration around 13ms. The mean interspike interval for all neurons in our sample is 12.7ms, corresponding to a rate of 78.7 spikes per second.

The functional bandwidth of transduction in vestibular hair cells and signal transmission in the vestibular nerve exceeds 1KHz (Bechstedt & Howard, 2007; Eatock, 2018; Hudspeth & Markin, 1994; Roberts, Howard, & Hudspeth, 1988). Such high bandwidth vestibular sense data must be ecologically important because otherwise evolution would not have continued to invest in molecular biophysical machinery capable of transducing it and delivering it to the brain. The prohibitive energy cost of firing at high rates provides strong selection pressure for mammals to distribute high bandwidth sensory signals over many neurons each firing at average rates in the order of tens of spikes per second (Balasubramanian, 2015; Sengupta & Stemmler, 2014).

The Exwald is the distribution of the sum of samples from Exponential and Wald distributions, but because the shape of an Exponential beyond any point is the same as the shape of the whole distribution, the Exwald is also the shape of an Exponential distribution left-censored by a Wald distribution. Thus the Wald component of an Exwald distribution can be interpreted as the distribution of refractory periods in a refractory-censored Poisson process. Refractory censoring with a mean interval around 12.5ms keeps average firing rate below 80s^−1^. Censoring with random rather than fixed refractory periods means that the censored samples are independent random samples from the uncensored distribution. Many such channels can transmit the same information in parallel at the same rate as a single neuron firing fast enough (i.e. sampling from the same distribution fast enough) to transmit the signal at high bandwidth, without incurring the catastrophic energy cost which that would entail. Thus the functional organization of the vestibular nerve is consilient with the proposal of Sterling and Laughlin (2015) that sensory neurons evolved to transmit information about molecular-scale transduction events at the mesoscale of whole animals, severely constrained by the energy costs of doing this using molecular mechanisms.

The heterogeneity of afferent neuron firing behaviour, i.e. the fact that they sample from Exwald distributions with different parameters, can be explained by the fact that distributed signalling on parallel channels can be more efficient if different channels have different characteristics (Barlow, 1961). As a specific example, it’s easier to detect larger signals in noise, and energy can be saved by using specialized channels with different sensitivities (Doi & Lewicki, 2014; van Hateren, 1992). This principle has been used to explain the statistical and dynamic diversity of retinal ganglion cells, and might also explain statistical and dynamical diversity among vestibular afferent neurons. The specific pattern of heterogeneity would depend on the statistics of natural head motion, which have not been well characterized in any species. Further investigation is required to explore and test whether the distribution of parameters illustrated in figure 5 reflects an optimally efficient way to distribute information about natural vestibular sense data across parallel channels, when individual channels are point processes with Exwald interval statistics.

Under general assumptions about spiking energetics, spike trains with Generalized Inverse Gaussian interval distributions maximize the information capacity of point-process channels subject to an energy constraint (Berger, Levy, & Jie, 2011; Xing, Berger, Sungkar, & Levy, 2015). Wald distributions are members of this class, suggesting that selection for energy efficiency may at least in part explain not only the massive parallelism and heterogeneity of information transmission in the vestibular nerve, but also the statistical distribution of intervals in individual neurons.

Natural selection does not act on components independently, but on the contribution of components to fitness of the organism. Thus the cost of information transmission in a sensory nerve must be weighed against the cost of processing that information in the brain. Other things being equal, we might expect brains to have evolved to be Bayesian (Levy, 2006), because Bayesian inference is 100% efficient in extracting information about parameters from data (Zellner, 1988)and is necessary for statistically optimal decision-making and optimal stochastic control (Berger, 1985). However, as critics of the “Bayesian brain” hypothesis have asked, at what cost? The mathematical efficiency of Bayesian inference does not account for the energy costs of Bayesian computation. Current Bayesian methods are computationally intensive, and indeed Bayesian inference has only recently become feasible beyond a few classical special cases as a result of massive reductions in the cost of computing (Kruschke, 2015). This suggests that even if were possible for neurons to extract information from sense data by Bayesian inference, the very high energy cost of neural computation should have weighed heavily against it, and should have favoured computationally cheaper heuristic rules and approximations instead (Bowers & Davis, 2012; Domurat, Kowalczuk, Idzikowska, Borzymowska, & Nowak-Przygodzka, 2015; Gigerenzer & Gaissmaier, 2011). Perhaps animals ought to behave as if they are Bayesians, as they do (McNamara, Green, & Olsson, 2006; Valone, 2006), but it is far from obvious that they should, could or actually do this by being Bayesian.

We found that Wald distributions by themselves are poor models, but convolution of Wald distributions with Exponential distributions produces Exwald distributions, which are excellent models of semicircular canal afferent neuron behaviour. The Exwald has an interesting property that may be relevant to Bayesian neural computation. A neuronal membrane compartment can be modelled electrically as a resistor parallel to a capacitor, whose response to impulsive current injection is an exponential decay function. Its response to an arbitrary current waveform is the convolution of that waveform with an exponential. Therefore, as we showed above, a membrane compartment containing a single synapse is a natural computer which can compute the likelihood of particular parameters given input pulses with Wald-like waveforms and Exwald-distributed intervals, instantaneously at all times. This property depends on having a series Poisson component in the data-generating process, and on synapses that can be parameterized so that the shape of EPSP matches the shape of the interval distribution of the other component. It is not necessary for the other component to have a Wald distribution. Thus while Bayesian computations have a reputation for being computationally intensive, slow and energy-hungry, there may be a simple, fast, efficient way for neurons to compute posterior densities of the parameters of a point process with Exwald interval distributions, given samples from the process. We speculate that Exwald distributions may have been selected for information transmission by vestibular afferent neurons because this optimizes energy efficiency, accounting for the total cost of data transmission and Bayesian inference for optimal dynamical head-state estimation in the brain.

Our goal was to construct a generative model of vestibular sense data, providing a foundation for developing testable models of neural computation for Bayesian inference in the vestibular system. We found that afferent spike trains are samples from stationary renewal processes with Exwald interval distributions. This model provides tantalizing hints about possible mechanisms of neural computation for Bayesian inference, and pointers for further research.

## Acknowledgements

The authors gratefully acknowledge the support of the NIH toward the conduct of this research (1R01DC014368 to LFH and MGP, and 1R01DC005059 to LFH).

